# Preoperative thymic output and T-cell repertoire diversity are associated with recurrence after nephrectomy for localized renal cell carcinoma

**DOI:** 10.64898/2026.10.13.757294

**Authors:** Christina Espeseth Grand, Laura Iisager, Asbjørn Kjær Attermann, Iver Nordentoft, Laura Zanini, Randi Istrup Juul, Anna Krarup Keller, Mie Gaedt Thorlund, Anne-Sofie Vibæk Eisum, Tommy Kjærgaard Nielsen, Jesper Jespersen, Niels Fristrup, Lars Dyrskjøt, Nicolai Juul Birkbak, Iben Lyskjær

**Affiliations:** Department of Molecular Medicine, Aarhus University Hospital, Aarhus, Denmark; Department of Clinical Medicine, Aarhus University, Aarhus, Denmark; Department of Urology, Aarhus University Hospital, Aarhus, Denmark; Department of Urology, Aalborg University Hospital, Aalborg, Denmark; Department of Urology, Odense University Hospital, Odense, Denmark; Department of Oncology, Aarhus University Hospital, Aarhus, Denmark; Department of Clinical Medicine, Aalborg University Hospital, Aalborg, Denmark

**Keywords:** Renal Cell Carcinoma, Thymic output, T-cell receptor repertoire, Peripheral blood, Prognosis, Leibovich score

## Abstract

Radiographic thymic health and peripheral T-cell repertoire features have been linked to outcomes in renal cell carcinoma (RCC), but their prognostic value after curative-intent surgery is unclear. We quantified preoperative blood signal-joint T-cell receptor excision circles (sjTRECs), a marker of thymic output, in 218 patients with localized RCC undergoing nephrectomy and characterized T-cell receptor β (TCRβ) repertoires in 201 patients. We assessed overall survival (OS) and recurrence-free survival (RFS) using Kaplan-Meier and Cox models adjusted for age, sex, and Leibovich risk group.

High sjTREC levels were associated with longer RFS after adjustment for age, sex, and Leibovich risk group (hazard ratio [HR] 0.50; 95% confidence interval [CI], 0.29-0.87; p=0.015), with a concordant OS estimate (HR 0.62; 95% CI, 0.38-1.02; p=0.058). sjTREC provided additional prognostic information beyond Leibovich risk stratification for recurrence overall and among intermediate/high-risk patients (likelihood-ratio test, p=0.024 and p=0.034, respectively). Higher TCRβ diversity was associated with longer RFS overall (HR 0.76; 95% CI, 0.60-0.96; p=0.024) and among intermediate/high-risk patients with longer RFS (HR 0.71; 95% CI, 0.55-0.92; p=0.009) and OS (HR 0.80; 95% CI, 0.65-0.98; p=0.031).

Preoperative sjTREC provided prognostic information beyond clinicopathological risk, with higher TCRβ diversity associated with favorable outcomes.

## Main text

The adult thymus is increasingly recognized as an important determinant of immune competence and clinical outcome. Preserved radiographic thymic health, assessed by the amount and composition of residual thymic tissue on imaging, has been associated with lower mortality in adults and improved outcomes following immune checkpoint inhibition. In operable lung cancer, greater thymic health was associated with longer disease-free survival, higher blood signal-joint T-cell receptor excision circle (sjTREC) levels and greater peripheral T-cell receptor (TCR) diversity, linking thymic health to both clinical outcome and circulating T-cell biology [1, 2]. sjTRECs are circular DNA by-products of T-cell receptor rearrangement and provide an indirect blood-based measure of thymic output, while TCR repertoire diversity reflects the breadth and clonal composition of the peripheral T-cell compartment. Peripheral-blood TCRβ repertoire diversity has also been associated with outcomes across several cancer types, including advanced RCC, although findings have varied across studies and treatment settings [3-5].

After nephrectomy for localized renal cell carcinoma (RCC), risk assessment is largely tumor-based, commonly using the Leibovich score [6]. Additional risk stratification is particularly relevant in intermediate/high-risk disease, where adjuvant treatment and intensified surveillance may be considered [7]. Whether host-related blood-based markers provide prognostic information remains unclear. We therefore investigated whether preoperative blood sjTREC levels and TCRβ repertoire diversity were associated with recurrence-free survival (RFS) and overall survival (OS), and whether they provided prognostic information beyond established clinicopathological risk.

We analyzed preoperative blood-derived DNA samples from 221 patients with localized RCC undergoing nephrectomy. sjTRECs were quantified in 218 patients, and TCRβ repertoires were characterized in 201 patients. For sjTREC analyses, patients were grouped according to the cohort median. TCRβ repertoire diversity was quantified using Gini-Simpson diversity and modeled continuously per 1-SD increase; normalized Shannon diversity was assessed in sensitivity analyses. Associations with RFS and OS were assessed using Kaplan-Meier analysis and Cox models adjusted for age, sex, and Leibovich risk group. Competing-risk analyses for recurrence applied Aalen-Johansen estimates and Fine-Gray models to account for death without recurrence. The additional prognostic value of sjTREC beyond Leibovich risk group was assessed using likelihood-ratio tests and bootstrap optimism-corrected C-indices. Cohort characteristics, follow-up, and event counts are summarized in Table 1.

**Table 1:** Cohort characteristics. Data are presented as median (Q1, Q3) or n (%). Abbreviations: Q1, first quartile; Q3, third quartile; sjTREC, signal-joint T-cell receptor excision circle; TCRβ, T-cell receptor β.

| **Characteristic** | **N = 221<sup>1</sup>** |
| --- | --- |
| **Age, yr** | 65 (56, 73) |
| **Sex** |  |
| Men | 141 (63.8%) |
| Women | 80 (36.2%) |
| **Histology** |  |
| Clear cell | 195 (88.2%) |
| Papillary | 21 (9.5%) |
| Chromophobe | 4 (1.8%) |
| Not recorded | 1 (0.5%) |
| **Tumor stage** |  |
| I | 90 (40.7%) |
| II | 31 (14.0%) |
| III | 96 (43.4%) |
| Localized, stage not specified | 4 (1.8%) |
| **Leibovich risk group** |  |
| Low | 62 (28.1%) |
| Intermediate | 87 (39.4%) |
| High | 72 (32.6%) |
| **Follow-up, yr** | 7.1 (3.8, 9.9) |
| **Recurrence** | 62 (28.1%) |
| **Death** | 75 (33.9%) |
| **sjTREC measurement available** | 218 (98.6%) |
| **TCRβ repertoire available** | 201 (91.0%) |

<sup>1</sup>Median (Q1, Q3); n (%)

Kaplan-Meier analysis showed longer RFS and OS in patients with high versus low sjTREC levels (log-rank p = 0.003 and p < 0.001, respectively, Fig. 1A+B). Aalen-Johansen cumulative incidence curves showed a similar separation between sjTREC groups (Fig. S1). After adjustment for age, sex, and Leibovich risk group, high sjTREC remained associated with longer RFS (HR 0.50; 95% CI, 0.29-0.87; p = 0.015, Fig. 1C and Fig. S2A), whereas the OS association was attenuated (HR 0.62; 95% CI, 0.38-1.02; p = 0.058, Fig. 1C and Fig. S2B). The association with RFS was also observed when sjTREC was analyzed continuously (Fig. S3). Among the clinically relevant intermediate/high-risk patients, high sjTREC was associated with longer RFS (HR 0.50; 95% CI, 0.28-0.88; p=0.017, Fig. S4). Adding sjTREC to the Leibovich model improved model fit in both the full cohort and the intermediate/high-risk subgroup (likelihood-ratio p = 0.024 and p = 0.034, respectively), with modest improvement in discrimination (Table S1).

**Figure 1:**
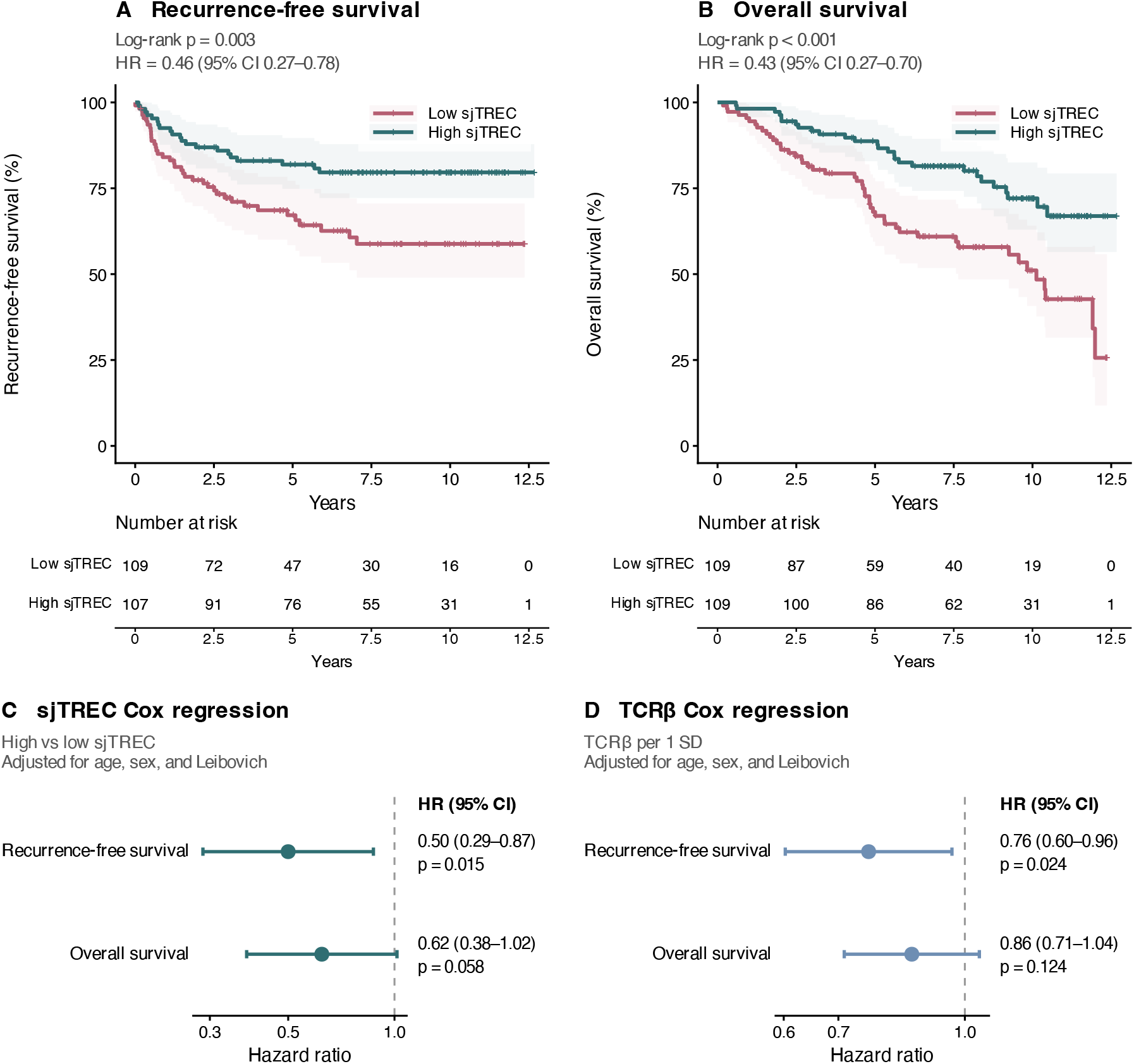
sjTREC levels and TCRβ diversity are associated with recurrence. **(A)** Kaplan-Meier analysis of RFS according to sjTREC level, dichotomized at the cohort median. (B) Kaplan-Meier analysis of OS according to sjTREC level, dichotomized at the cohort median. (C) Forest plot showing Cox regression analyses of RFS and OS according to sjTREC group adjusted for age, sex, and Leibovich risk group. (D) Forest plot showing Cox regression of RFS and OS according to continuous Gini-Simpson TCRβ diversity, adjusted for age, sex, and Leibovich risk group. HRs are reported per 1-SD increase in diversity. For the Kaplan-Meier panels, p-values were obtained using the log-rank test, and HRs were estimated using unadjusted Cox models. Shaded areas represent 95% confidence intervals. Abbreviations: CI, confidence interval; HR, hazard ratio; OS, overall survival; sjTREC, signal-joint T-cell receptor excision circle; TCRβ, T-cell receptor β; RFS, recurrence-free survival; SD, standard deviation.

We also assessed peripheral TCRβ diversity, which correlated only weakly with sjTREC levels (Spearman ρ = 0.20; p = 0.004, Fig. S5). Higher TCRβ diversity was associated with longer RFS in the full cohort after adjustment for age, sex, and Leibovich risk group (HR 0.76; 95% CI, 0.60-0.96; p = 0.024; Fig. 1D and S2C), while the OS estimate was directionally similar (HR 0.86; 95% CI, 0.71-1.04; p=0.124; Fig. 1D and S2D). Among intermediate/high-risk patients, higher TCRβ diversity was associated with longer RFS (HR 0.71; 95% CI, 0.55-0.92; p=0.009, Fig. S6C) and OS (HR 0.80; 95% CI, 0.65-0.98; p=0.031, Fig. S6D). Normalized Shannon diversity showed associations in the same direction, although none reached statistical significance in either population (Fig. S6).

Adjusted competing-risk analyses treating death without prior recurrence as a competing event yielded similar associations for sjTREC and TCRβ diversity (Table S2). Associations with both sjTREC and TCRβ diversity were also observed in analyses restricted to ccRCC, the histological subtype for which the Leibovich score was originally developed (Fig. S7). When sjTREC and TCRβ diversity were included in the same multivariable model, their effect estimates were largely preserved, suggesting that their associations with outcome were not explained by each other (Fig. S8).

We found that in patients undergoing nephrectomy for localized RCC, higher preoperative sjTREC level was associated with longer RFS after adjustment for age, sex, and Leibovich risk group, with a concordant but nonsignificant OS estimate. sjTREC also provided additional prognostic information for recurrence beyond the Leibovich risk group, including among intermediate/high-risk patients. These findings extend previous observations linking radiographic thymic health to clinical outcome [2] by showing that a blood-based marker of thymic output carries prognostic information before curative-intent surgery. Importantly, sjTREC can be quantified using a relatively simple and inexpensive droplet digital PCR assay, supporting its potential for clinical implementation. TCRβ diversity provided a complementary signal: higher diversity was associated with longer RFS overall and with both longer RFS and OS among intermediate/high-risk patients. The weak correlation between sjTREC and TCRβ diversity and the preservation of their effect estimates in a joint model suggest that they capture partly distinct aspects of systemic T-cell state.

The study has several limitations. This was a retrospective, exploratory study, and the findings were not evaluated in an independent validation cohort. Peripheral sjTREC is an indirect marker of thymic output and may also be influenced by peripheral T-cell proliferation, and the optimal analytical threshold remains unknown. Prospective validation using a predefined assay and threshold is required.

In summary, preoperative sjTREC provided prognostic information for recurrence beyond established clinicopathological risk, while TCRβ diversity showed a complementary association with outcomes. These findings support prospective evaluation of systemic T-cell measures for postoperative risk stratification.

## Declarations

### Ethics

approved by the Danish National Center for Ethics (protocol 2208138).

## Acknowledgements

We thank the Bio- and Genome Bank Denmark and the Danish Cancer Biobank for provision, handling and storage of biological material and associated data, GenomeDK for computing resources, and the patients who donated material. We thank lab technicians Lotte Gernyx, Lisbet Kjeldsen, Helena Ellegaard Haahr, Margaret Elizabeth Gellett, and Gitte Glistrup Nielsen for handling our samples.

## Availability of data and materials

Data are available through controlled access from GenomeDK (https://genome.au.dk/library/GDK000024/). Code is available on GitHub https://github.com/Christinagrand/sjTREC_TCRB.

## Author contributions

Iben Lyskjær had full access to all study data and is responsible for data integrity and the accuracy of the data analysis. **Study concept and design**: Grand, Iisager, Birkbak, Dyrskjøt, Lyskjær. **Acquisition of data**: Grand, Iisager, Attermann, Zanini, Nordentoft, Keller, Thorlund, Eisum, Nielsen, Fristrup, Lyskjær. **Analysis and interpretation of data:** Grand, Iisager, Attermann, Zanini, Juul, Jespersen, Birkbak, Lyskjær. **Statistical analysis:** Grand. **Drafting of the manuscript:** Grand, Iisager, Attermann, Lyskjær. **Critical revision of the manuscript for important intellectual content:** all authors. **Obtaining funding:** Iisager, Lyskjær. **Supervision**: Lyskjær.

## Declaration of Generative AI and AI-assisted technologies in the writing process

During preparation of this manuscript, the authors used a large language model to assist with language editing and condensation, and to assist with writing and reviewing analysis code. All code was reviewed and executed by the authors. The authors reviewed and edited any AI-assisted content and take full responsibility for the content of the publication.

