## Supplementary method, figures and tables for "Preoperative thymic output and T-cell repertoire diversity are associated with recurrence after nephrectomy for localized renal cell carcinoma"

**Supplementary Methods**

**Patients and cohorts**
Patients underwent partial or radical nephrectomy for localized RCC (stages I-III) between 2010 and 2021 and had a preoperative peripheral blood sample available through the Danish Cancer Biobank at Bio- and Genome Bank Denmark (RBGB). Clinical data were obtained from RBGB and the Danish Clinical Quality Program (RKKP). The patient cohort was selected based on availability of clinical data. Clinical follow-up included recurrence status, time to recurrence, vital status, and time to death, obtained from the RKKP databases and, where necessary, from medical records. Cause-of-death data were not available. Patients with missing recurrence dates were excluded from recurrence analyses (two in the sjTREC cohort, three in the TCRβ cohort).

sjTREC data were generated for 251 patients, of whom 32 were excluded due to missing Leibovich classification, and one was excluded as an extreme outlier, with an sjTREC value approximately 494-fold higher than the second-highest observation. The final sjTREC analysis cohort comprised 218 patients, with cohort characteristics summarized in Table 1.

TCRβ repertoire data were generated for 251 patients. Of these, 16 patients were excluded due to failed quality control, eight due to fewer than 500 unique productive TCRβ clonotypes, and 26 due to missing Leibovich classification. The final TCRβ analysis cohort comprised 201 patients.

**Sample collection and DNA extraction**
Buffy coat samples (50-4500 µL based on availability) were subjected to DNA extraction using the QIAsymphony DSP DNA Midi kit (QIAGEN) on a QIAsymphony SP instrument. DNA concentration was determined by Qubit Fluorometric (ThermoFisher Scientific) and DropSense96 (Trinean).

**sjTREC digital droplet PCR**
sjTREC levels were quantified by multiplex droplet digital PCR (ddPCR) using the QX200 AutoDG ddPCR System (Bio-Rad), as previously described [1]. sjTREC abundance was normalized to the diploid reference gene *RPP30*. For each sample, 600 ng genomic DNA was analyzed per reaction in triplicate (1,800 ng total). DNA was pretreated with HindIII restriction enzyme (BioNordika) for linearization (20 min at 37 °C followed by 10 min at 80 °C).

Reactions were performed using 2X ddPCR Supermix for Probes (no dUTP; Bio-Rad) with custom primer-probe sets targeting sjTREC and *RPP30*. The sjTREC primers were 5′-CACATCCCTTTCAACCATGCT-3′ (forward) and 5′-GCCAGCTGCAGGGTTTAGG-3′ (reverse), with probe 5′-ACACCTCTGTTTTTGTAAAGGTGCCCACT-3′ (6-FAM, Bio-Rad). The *RPP30* primers were *5′-AGATTTGGACCTGCGAGCG-3′ (forward)* and *5′-GAGCGGCTGTCTCCACAAGT-3′ (reverse)* with probe *5′-TTCTGACCTGAAGGCTCTGCGCG-3′* (HEX, Bio-Rad), as previously published [1].

Nuclease-free water was included as a no-template control. A mix of buffy-coat DNA from eight young healthy volunteers (younger than 30 years) was used as a positive control for *RPP30* and sjTREC, and a synthetic sjTREC gBlock fragment containing the primer target was used as a positive control for sjTREC detection. Data were analyzed using QX Manager software (Bio-Rad). sjTREC abundance was calculated as: ‘sjTREC per cell = 2 · FAM concentration (copies/μL) / HEX concentration (copies/μL)’, where the factor of 2 accounts for the diploid *RPP30* reference gene. The mean of the three replicate measurements was used as the final sjTREC value for each sample. If RPP30 was not detected in one replicate, the mean of the two remaining replicates was used. The distribution of sjTREC values is shown in S5A.

**TCRβ sequencing
Library preparation and sequencing**
*TCRβ* sequencing libraries were prepared using the Cellecta DriverMap AIR TCR Profiling Kit (Cellecta, Mountain View, CA, USA; Human DNA; version 1 and 2) with minor modifications. For each sample, 3,000-5,000 ng of genomic DNA was used as input. Libraries were amplified using 22-25 PCR cycles, depending on DNA input, followed by bead-based purification and elution in 30 µL of EB buffer. Library quality was assessed for each sample using TapeStation4200 (Agilent) before pooling. Libraries were subsequently paired-end sequenced with 148 bp reads on an Illumina NovaSeq6000 platform using S1, S2, and S4 flow cells. DNA concentration was determined by Qubit Fluorometric (ThermoFisher Scientific) and DropSense96 (Trinean).

DriverMap AIR kit version differed between patients. The first 140 patient samples were prepared with version 1, which incorporates random unique molecular identifiers (UMIs), whereas the next 111 patient samples were prepared with version 2, which uses validator barcodes for template-molecule quantification. Consequently, the two kit versions differ in the number of clonotypes recovered. TCRβ diversity was comparable between kit versions and diversity measures were therefore pooled across versions (Fig. S9).

For version 1, sequencing yielded a median of 18.8 million TCRβ-assigned reads (IQR, 13.7-24.5 million) before filtering of non-productive clonotypes. After filtering, a median of 5,134 template molecules (IQR, 3,016-8,200) and 3,556 unique TCRβ clonotypes (IQR, 2,114-5,364) were retained per patient. Corresponding values for version 2 were 4.0 million TCRβ-assigned reads (IQR, 2.9-5.0 million), 12,441 template molecules (IQR, 9,913-19,329), and 7,899 unique TCRβ clonotypes (IQR, 5,973-11,440). Table S3 provides sequencing metrics by kit version.

**Repertoire reconstruction and quality filters**
Base call files were converted to FASTQ and demultiplexed by index sequence using Illumina bcl2fastq (v2.20.0). Repertoire reconstruction was performed using MiXCR v4.7.0 (RepSeq.IO v5.1) with the cellecta-human-dna-xcr-umi-drivermap-air preset for samples sequenced with the first version of the kit, and MiXCR v4.7.0-309-develop with the cellecta-human-dna-xcr-mivbc-drivermap-air-v2 preset for samples sequenced with the second version of the kit [2, 3]. For the second version of the kit, MiXCR output was subsequently processed using Cellecta’s VBC filtering and template-molecule estimation pipeline [4].

Samples with fewer than 500 unique productive TCRβ clonotypes were excluded because repertoire recovery below this threshold was insufficient for robust diversity estimation.

Clone abundances were calculated from molecule counts rather than read counts to avoid PCR amplification bias. Clonotypes with frameshifts and stop codons were excluded, as these rearrangements cannot encode a functional receptor. Clonotype frequencies were then recalculated within each sample by dividing the molecule count of each retained clonotype by the total molecule count of all retained clonotypes, ensuring that frequencies summed to one before calculating diversity indices. The same reconstruction and filtering procedures were applied to both kit versions.

Repertoires were reconstructed for all T-cell receptor chains. TCRβ was selected as the primary chain, because it is a conventional target for repertoire diversity analyses and the chain with the highest recovery of unique productive clonotypes in kit version 1.

**Diversity indices**
Two indices were computed from the clonotype frequencies of each sample, where $p_{i}$ is the frequency of clonotype i and S is the number of unique productive clonotypes in that sample.

Gini-Simpson diversity was the primary index: $D=1-\sum_{i=1}^{S} p_{i}^{2}$

It ranges from 0 to 1, with values near 0 indicating a repertoire dominated by few clonotypes and values near 1 indicating a repertoire in which clonotype frequencies are evenly distributed [5].

Normalized Shannon diversity was computed as a sensitivity analysis: $H_{norm}= \frac{-\sum_{i=1}^{S} p_{i}*ln(p_{i})}{ln(S)}$

**Standardization and scale**
Each diversity index was standardized using its mean and standard deviation across all patients with an available measurement in the final pooled cohort. The same standardization was retained in subgroup analyses. Hazard ratios therefore represent the effect of a one-standard-deviation increase in diversity.

**Visualization**
For visualization, TCRβ Gini-Simpson diversity was transformed as −log_10_(1 − Gini-Simpson) in Fig. S5C, D, and F, while sjTREC was displayed on a log_2_-transformed scale in Fig. S5A, D, and E. Axis labels show the original values. Group comparisons and correlations were assessed using untransformed values.

**Clinical variables and endpoints**
Overall survival (OS) was defined as the time from diagnosis to death from any cause. Recurrence-free survival (RFS) was defined as the time from diagnosis to first documented recurrence. Patients without recurrence were censored at death or the end of follow-up. Blood samples were collected preoperatively, shortly before nephrectomy.

**Statistical analysis**
Spearman’s rank correlation was used to assess associations between age and each immune measure, and between sjTREC levels and TCRβ Gini-Simpson diversity.

Kaplan-Meier analyses were performed with sjTREC dichotomized at the median of the full sjTREC analysis cohort, and groups were compared using the log-rank test. Values above the median were classified as high, and values at or below the median as low. The same threshold was retained in subgroup analyses. Cox proportional hazards models were fitted with TCRβ diversity as a continuous variable and sjTREC level as a dichotomous variable. Models were adjusted for age, sex, and Leibovich risk group. In sensitivity analyses, sjTREC was modeled continuously as log_2_(sjTREC + c), where c was half the smallest positive sjTREC value in the full analysis cohort. For continuous sjTREC analyses, HRs represent a one-unit increase in this transformed measure.

To assess the additional prognostic value of sjTREC for recurrence beyond the Leibovich risk group, likelihood-ratio tests were used to compare nested Cox models and assess whether adding sjTREC improved model fit beyond the Leibovich risk group. Discrimination was assessed using Harrell’s concordance index (C-index). The discriminative performance of a Cox model including Leibovich risk group alone was compared with that of a model additionally including dichotomized sjTREC levels. C-indices were internally validated using 2,000 bootstrap resamples to estimate and correct for optimism. Optimism-corrected C-indices were calculated by subtracting the mean estimated optimism from the apparent C-index. Analyses were performed in the full cohort and separately among patients with intermediate/high Leibovich-risk disease.

For competing-risk analyses of recurrence, death without prior recurrence was treated as a competing event. Cumulative incidence of recurrence was estimated using the Aalen-Johansen estimator, and differences between sjTREC groups were assessed using Gray's test. Multivariable Fine-Gray competing-risk regression was performed with adjustment for age, sex, and Leibovich risk group.

**Software**
Analyses were performed in R (v4.6.0) using the survival (v3.8-6), survminer (v0.5.2), tidycmprsk (v1.1.2), cmprsk (v2.2-12), ggsurvfit (v1.2.1), ggplot2 (v4.0.3), broom (v1.0.13), and patchwork (v1.3.2) packages. Statistical significance was defined as p < 0.05. All reported p-values are nominal.

**Supplementary Tables**

**Table S1. C-index for recurrence: Leibovich risk group alone and with sjTREC.** Corrected C-index was calculated as the apparent C minus the mean optimism from 2,000 bootstrap resamples. ΔC represents the corrected C of each model minus the corrected C of the Leibovich-only model in the same population. Likelihood ratio test (LRT) p-values were obtained from likelihood ratio tests comparing the model including sjTREC with the Leibovich-only model. sjTREC was dichotomized at the median of the full sjTREC analysis cohort, and the same cut-off was applied in both populations. Abbreviations: C-index, concordance index; LRT, likelihood ratio test; sjTREC, signal-joint T-cell receptor excision circle.

| **Model** | **Apparent C** | **Optimism** | **Corrected C** | **ΔC vs Leibovich** | **LRT p** |
| --- | --- | --- | --- | --- | --- |
| **All patients (n = 216; 59 recurrences)** | | | | | |
| Leibovich only | 0.7447 | 0.0002 | 0.7444 | Reference | — |
| Leibovich + sjTREC | 0.7695 | 0.0038 | 0.7657 | +0.0213 | 0.024 |
| **Intermediate/high Leibovich risk (n = 154; 55 recurrences)** | | | | | |
| Leibovich only | 0.6675 | -0.0001 | 0.6676 | Reference | — |
| Leibovich + sjTREC | 0.7035 | 0.0049 | 0.6987 | +0.0310 | 0.034 |

**Table S2. Fine-Gray competing-risk sensitivity analyses for recurrence.** Fine-Gray regression was performed with recurrence as the event of interest and death without prior recurrence as the competing event. Models were adjusted for age, sex, and Leibovich risk group. sjTREC levels were dichotomized at the cohort median. TCRβ Gini-Simpson diversity was modeled continuously per 1-SD increase. Abbreviations: CI, confidence interval; SD, standard deviation; SHR, subdistribution hazard ratio; sjTREC, signal-joint T-cell receptor excision circle; TCRβ, T-cell receptor β.

| **Variable** | **SHR** | **95% CI** | **p value** |
| --- | --- | --- | --- |
| **sjTREC model (n = 216; 59 recurrences; 35 competing deaths)** | | | |
| High vs low sjTREC | 0.51 | 0.29-0.89 | 0.017 |
| Age, per year | 0.98 | 0.96-1.01 | 0.160 |
| Sex: woman vs man | 0.75 | 0.40-1.38 | 0.350 |
| Leibovich: intermediate vs low | 3.40 | 1.13-10.19 | 0.029 |
| Leibovich: high vs low | 12.48 | 4.35-35.80 | <0.001 |
| **TCRβ model (n = 198; 53 recurrences; 32 competing deaths)** | | | |
| TCRβ diversity per 1-SD | 0.76 | 0.64-0.89 | 0.001 |
| Age, per year | 0.99 | 0.96-1.01 | 0.370 |
| Sex: woman vs man | 0.68 | 0.36-1.32 | 0.260 |
| Leibovich: intermediate vs low | 4.21 | 1.08-16.32 | 0.038 |
| Leibovich: high vs low | 17.85 | 5.07-62.85 | <0.001 |

**Table S3. TCRβ sequencing metrics by DriverMap AIR kit version.** Sequencing depth, productive template molecules, and unique productive TCRβ clonotypes are summarized separately for DriverMap AIR versions 1 and 2. Values are presented as median (IQR) and range. Abbreviations: IQR, interquartile range; TCRβ, T-cell receptor β.

| **Metric** | **Kit version** | **Median (IQR)** | **Range** |
| --- | --- | --- | --- |
| TCRβ-assigned reads | Version 1 | 18.8 (13.7-24.5) million | 4.4-76.8 million |
|  | Version 2 | 4.0 (2.9-5.0) million | 1.5-20.6 million |
| Productive template molecules | Version 1 | 5,134 (3,016-8,200) | 534-24,936 |
|  | Version 2 | 12,441 (9,913-19,329) | 785-30,646 |
| Unique productive TCRβ clonotypes | Version 1 | 3,556 (2,114-5,364) | 508-20,616 |
|  | Version 2 | 7,899 (5,973-11,440) | 762-23,852 |

**Supplementary Figures**

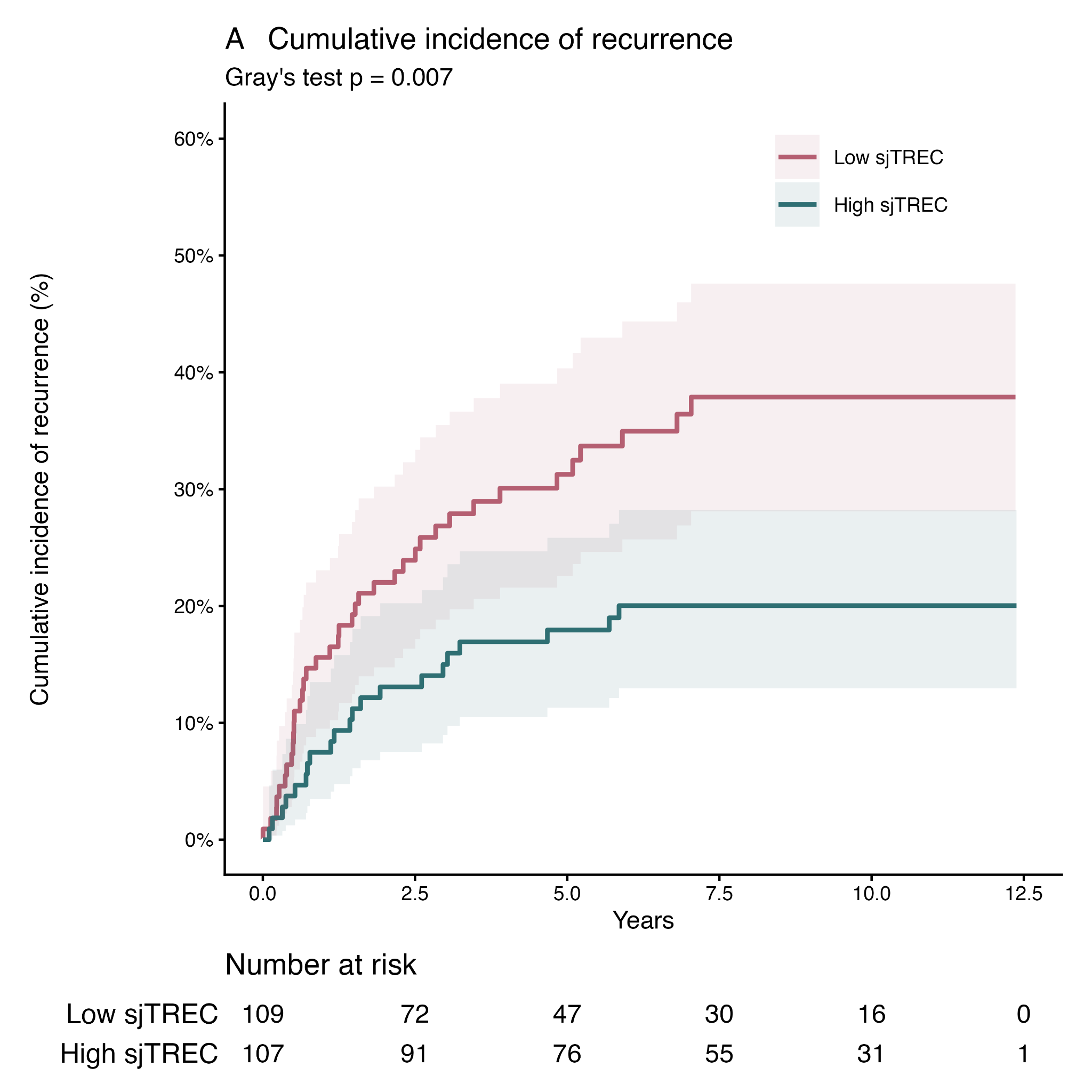

**Figure S1: Cumulative incidence of recurrence according to sjTREC levels.** Aalen-Johansen cumulative incidence curves for patients with low and high sjTREC levels, dichotomized at the cohort median. Death without prior recurrence was treated as a competing event. Shaded areas represent 95% confidence intervals. Differences between groups were assessed using Gray's test. Abbreviation: sjTREC, signal-joint T-cell receptor excision circle.

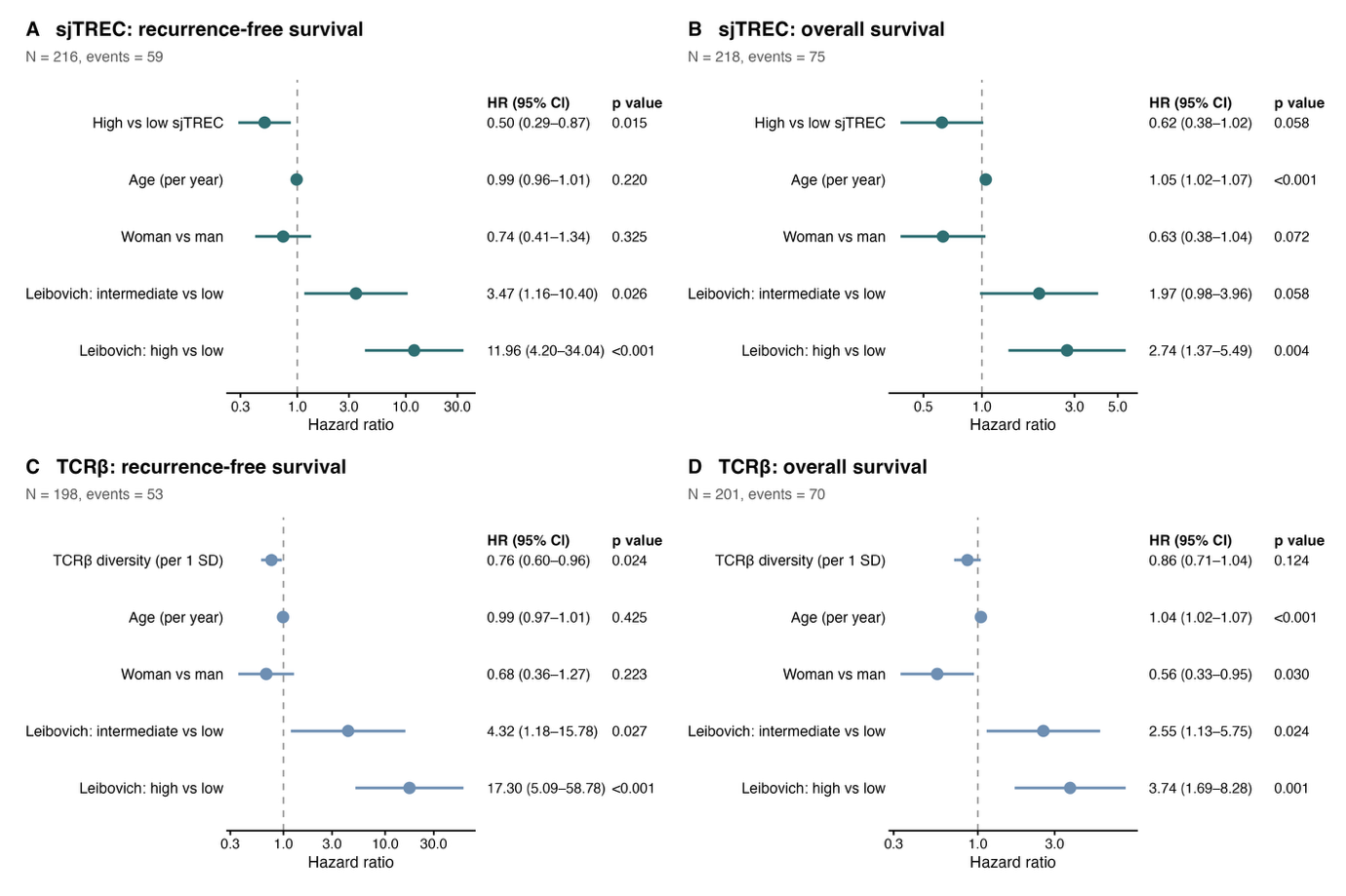

**Figure S2. Full forest plots of Cox regression models for dichotomized sjTREC and continuous TCRβ Gini-Simpson diversity.** sjTREC was dichotomized at the cohort median. (A) sjTREC for recurrence-free survival, (B) sjTREC for overall survival, (C) TCRβ Gini-Simpson diversity for recurrence-free survival, and (D) TCRβ Gini-Simpson diversity for overall survival. TCRβ diversity HRs are reported per 1-SD increase. Abbreviations: CI, confidence interval; HR, hazard ratio; SD, standard deviation; sjTREC, signal-joint T-cell receptor excision circle; TCRβ, T-cell receptor β.

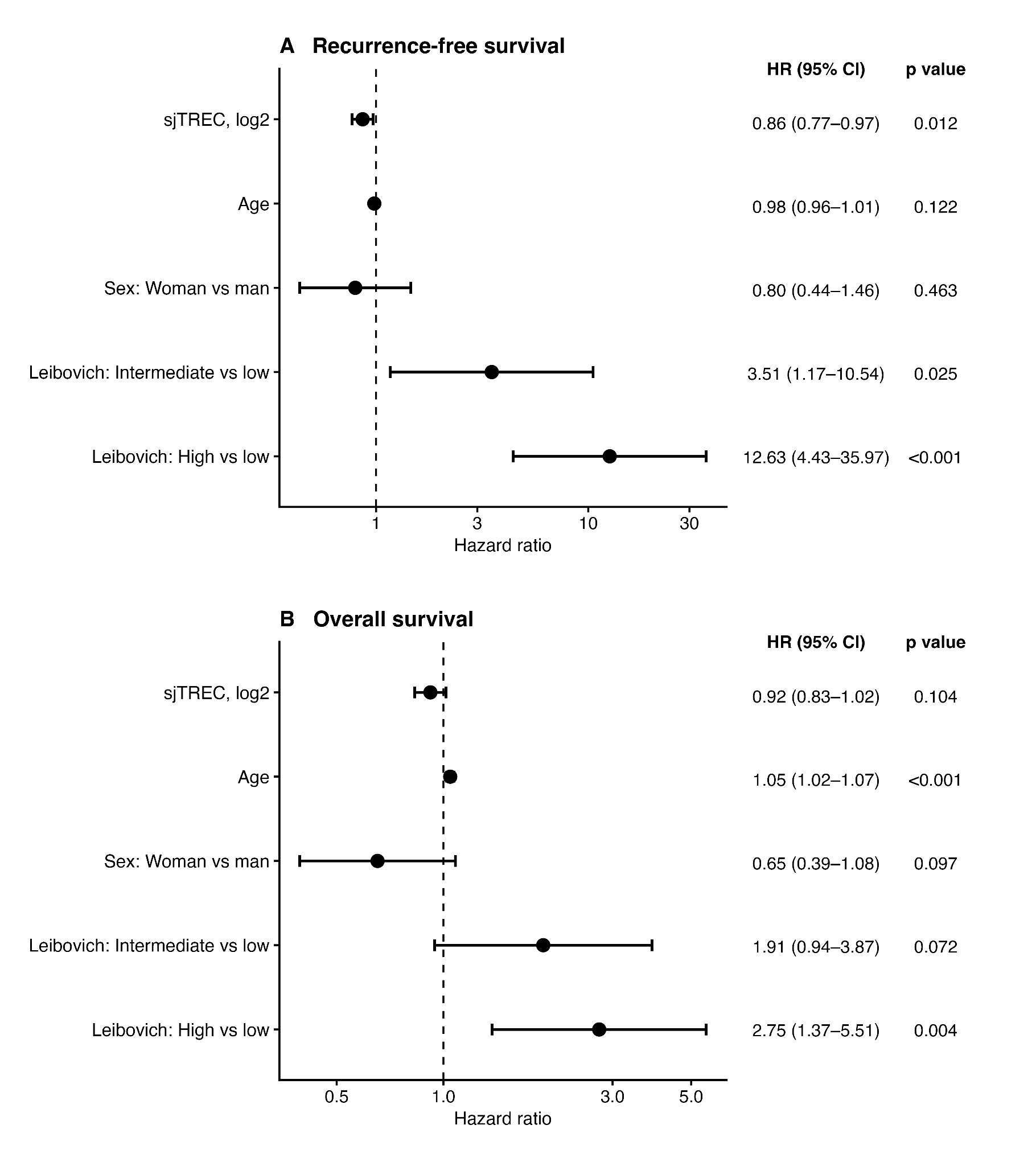

**Figure S3. Associations of continuous sjTREC with clinical outcomes.** Cox regression analyses of (A) recurrence-free survival and (B) overall survival with sjTREC modeled continuously on a log_2_-transformed scale. HRs represent a one-unit increase in the transformed measure. Cox models were adjusted for age, sex, and Leibovich risk group. Abbreviations: CI, confidence interval; HR, hazard ratio; sjTREC, signal-joint T-cell receptor excision circle.

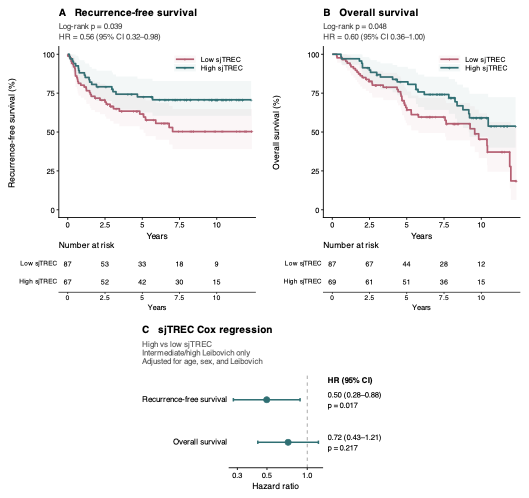

**Figure S4. Association of sjTREC with outcomes in intermediate/high-risk patients.** sjTREC was dichotomized at the median of the full sjTREC analysis cohort. (A) Kaplan-Meier analysis of recurrence-free survival according to sjTREC group, (B) Kaplan-Meier analysis of overall survival according to sjTREC group, and (C) Cox regression analyses of recurrence-free survival and overall survival. Cox models were adjusted for age, sex and Leibovich risk group; adjustment covariates are not shown. For the Kaplan-Meier panels, p-values were obtained using the log-rank test, and HRs were estimated using unadjusted Cox models. Shaded areas represent 95% confidence intervals. Abbreviations: CI, confidence interval; HR, hazard ratio; OS, overall survival; sjTREC, signal-joint T-cell receptor excision circle; RFS, recurrence-free survival.

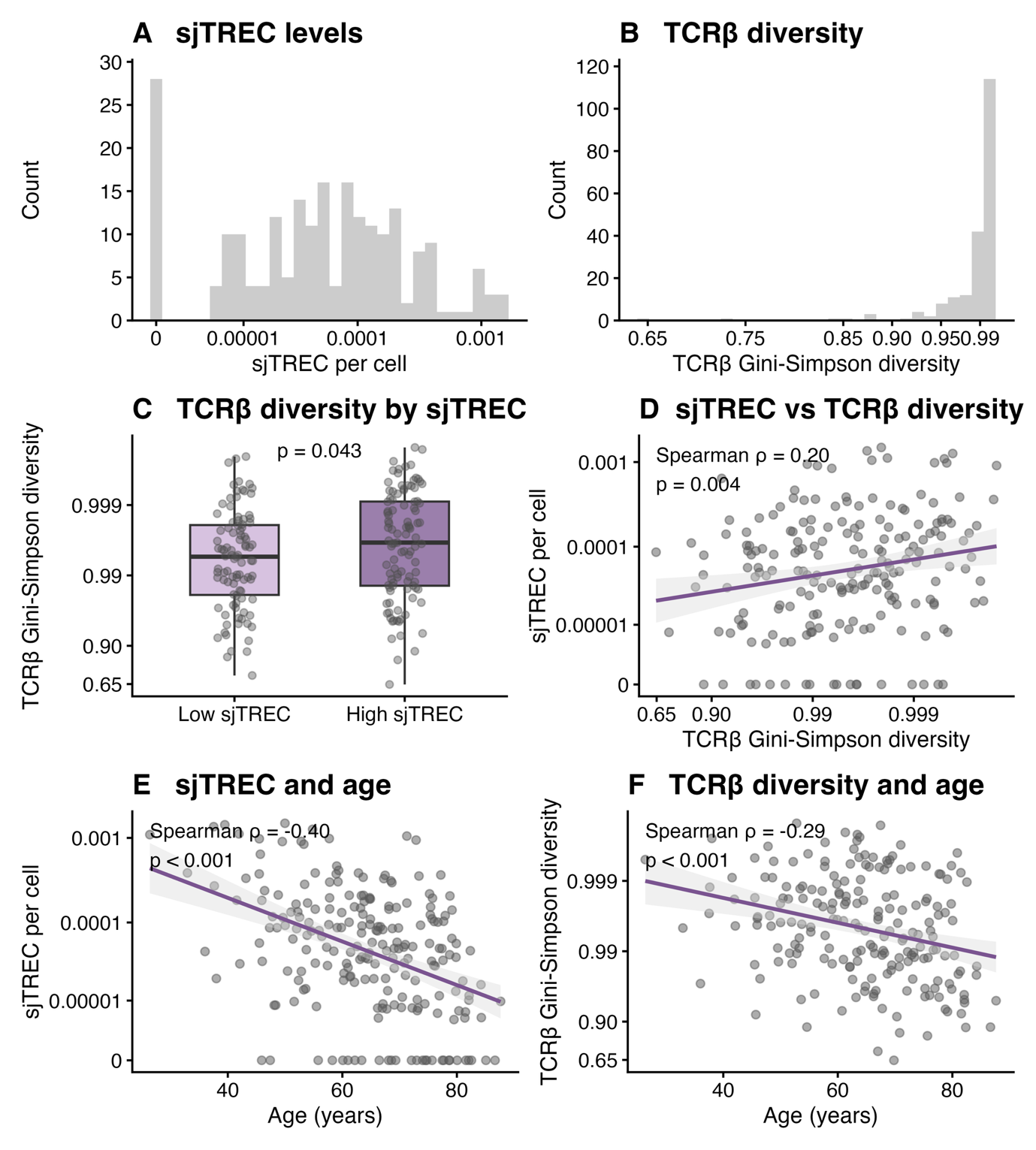

**Figure S5. Overview of sjTREC levels and TCRβ diversity.** (A) Distribution of sjTREC levels. (B) Distribution of TCRβ Gini-Simpson diversity. (C) TCRβ Gini-Simpson diversity according to low and high sjTREC groups, defined by the cohort median. (D) Correlation between sjTREC levels and TCRβ Gini-Simpson diversity. (E) Correlation between sjTREC levels and age. (F) Correlation between TCRβ Gini-Simpson diversity and age. Differences in TCRβ Gini-Simpson diversity between sjTREC groups were assessed using the Wilcoxon rank-sum test. Correlations were assessed using Spearman’s rank correlation. For visualization, sjTREC was displayed on a log_2_-transformed scale in panels A, D and E, and TCRβ Gini-Simpson diversity on a −log_10_(1 − Gini-Simpson) scale in panels C, D and F. Axis labels show the original values. Abbreviations: sjTREC, signal-joint T-cell receptor excision circle; TCRβ, T-cell receptor β.

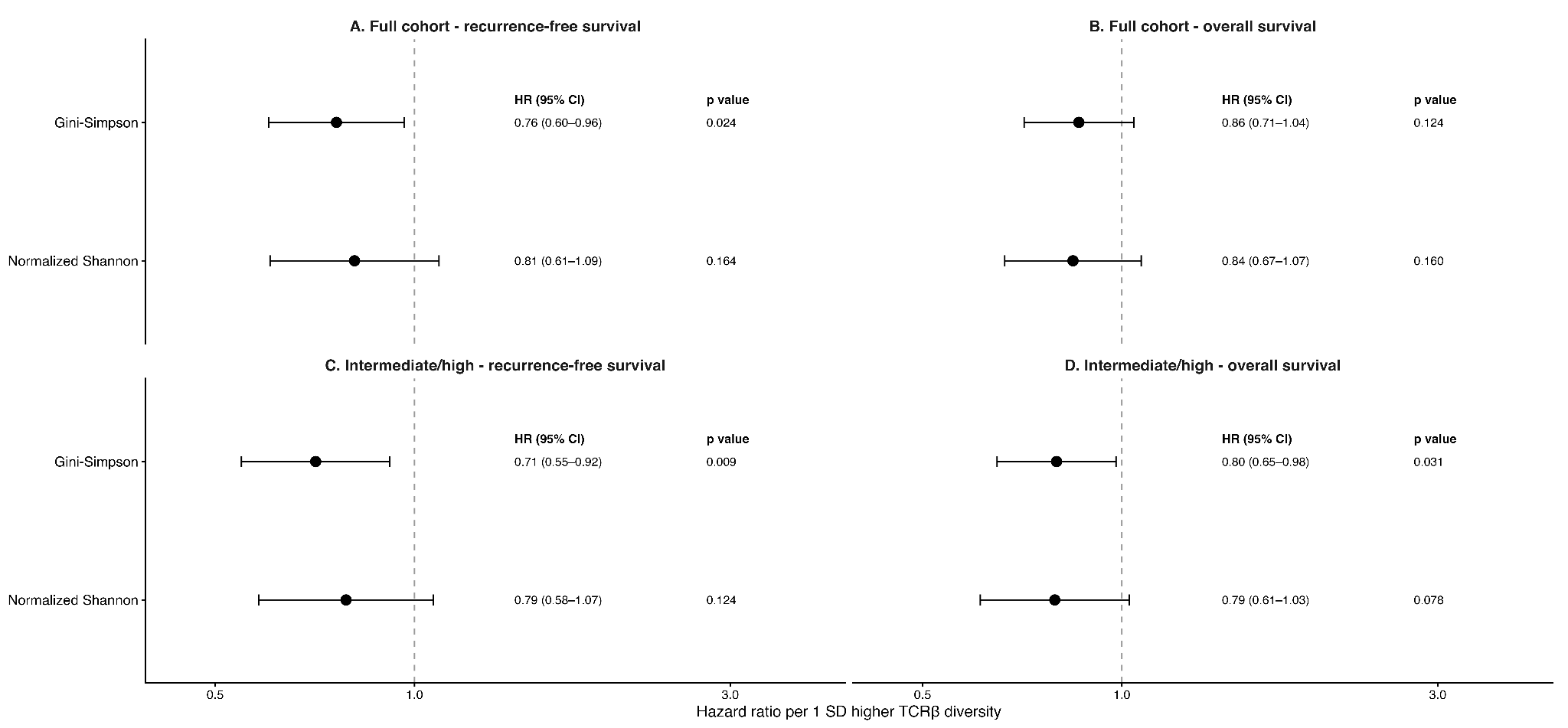

**Figure S6. Associations between TCRβ diversity and clinical outcomes.** Forest plots showing associations of TCRβ diversity, quantified using Gini-Simpson diversity and normalized Shannon diversity, with (A) recurrence-free survival in the full cohort, (B) overall survival in the full cohort, (C) recurrence-free survival in the intermediate/high Leibovich-risk subgroup, and (D) overall survival in the intermediate/high Leibovich-risk subgroup. Hazard ratios are reported per 1-SD increase in each diversity measure. All Cox models were adjusted for age, sex, and Leibovich risk group. Abbreviations: OS, overall survival; SD, standard deviation; TCRβ, T-cell receptor β; RFS, recurrence-free survival; CI, confidence interval; HR, hazard ratio.

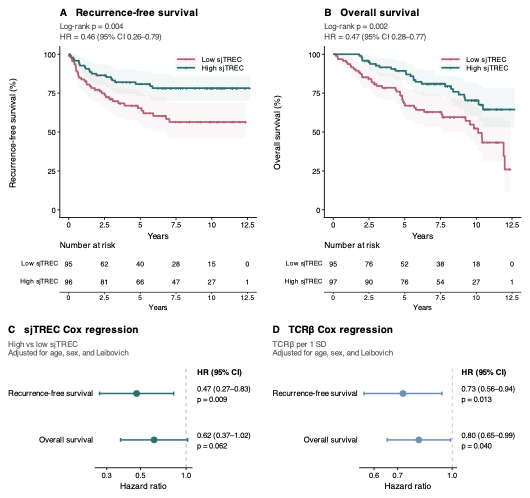

**Figure S7. Associations of sjTREC levels and TCRβ diversity with outcomes in patients with clear cell RCC.** (A) Kaplan-Meier analysis of RFS according to sjTREC level, dichotomized at the median of the full sjTREC analysis cohort. (B) Kaplan-Meier analysis of OS according to sjTREC level, dichotomized at the median of the full sjTREC analysis cohort. (C) Forest plot showing Cox regression analyses of RFS and OS according to sjTREC group, adjusted for age, sex and Leibovich risk group. (D) Forest plot showing Cox regressions of RFS and OS according to continuous TCRβ Gini-Simpson diversity, adjusted for age, sex and Leibovich risk group. HRs are reported per 1-SD increase in diversity. For the Kaplan-Meier panels, p-values were obtained using the log-rank test, and HRs were estimated using unadjusted Cox models. Shaded areas represent 95% confidence intervals. Abbreviations: CI, confidence interval; HR, hazard ratio; OS, overall survival; sjTREC, signal-joint T-cell receptor excision circle; TCRβ, T-cell receptor β; RFS, recurrence-free survival; SD, standard deviation.

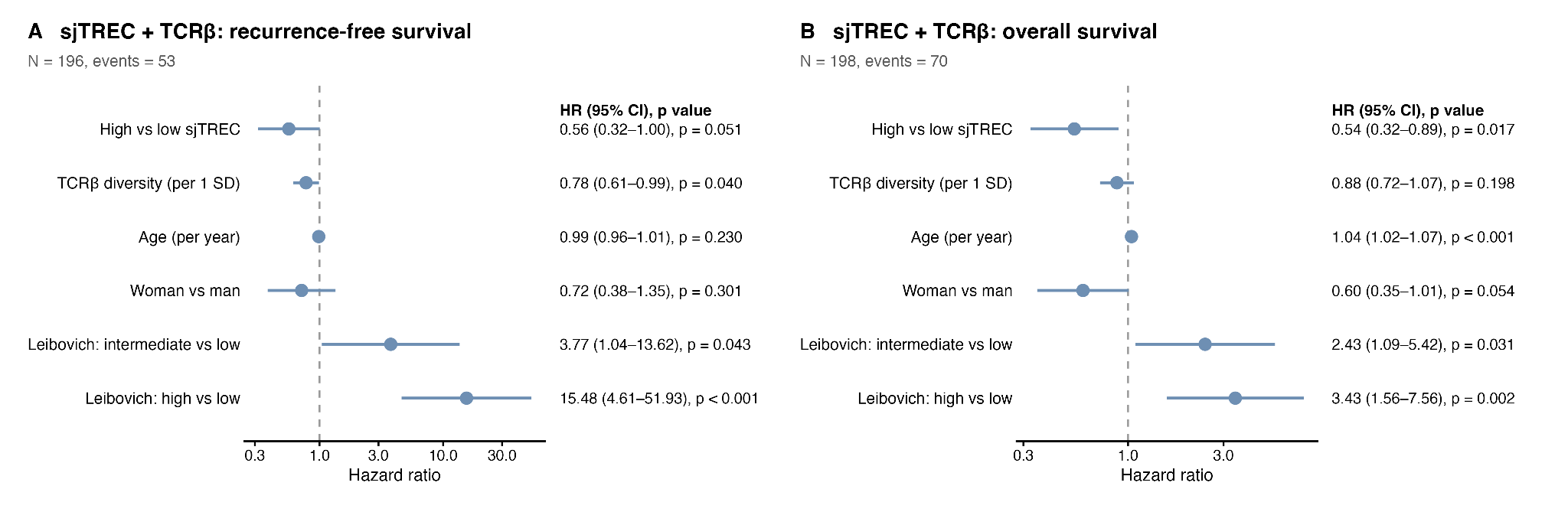

**Figure S8. Multivariable Cox regression including continuous TCRβ Gini-Simpson diversity and dichotomized sjTREC levels.** Analyses included patients with both immune measures available. sjTREC was dichotomized at the cohort median. (A) Recurrence-free survival. (B) Overall survival. Abbreviations: CI, confidence interval; HR, hazard ratio; sjTREC, signal-joint T-cell receptor excision circle; TCRβ, T-cell receptor β.

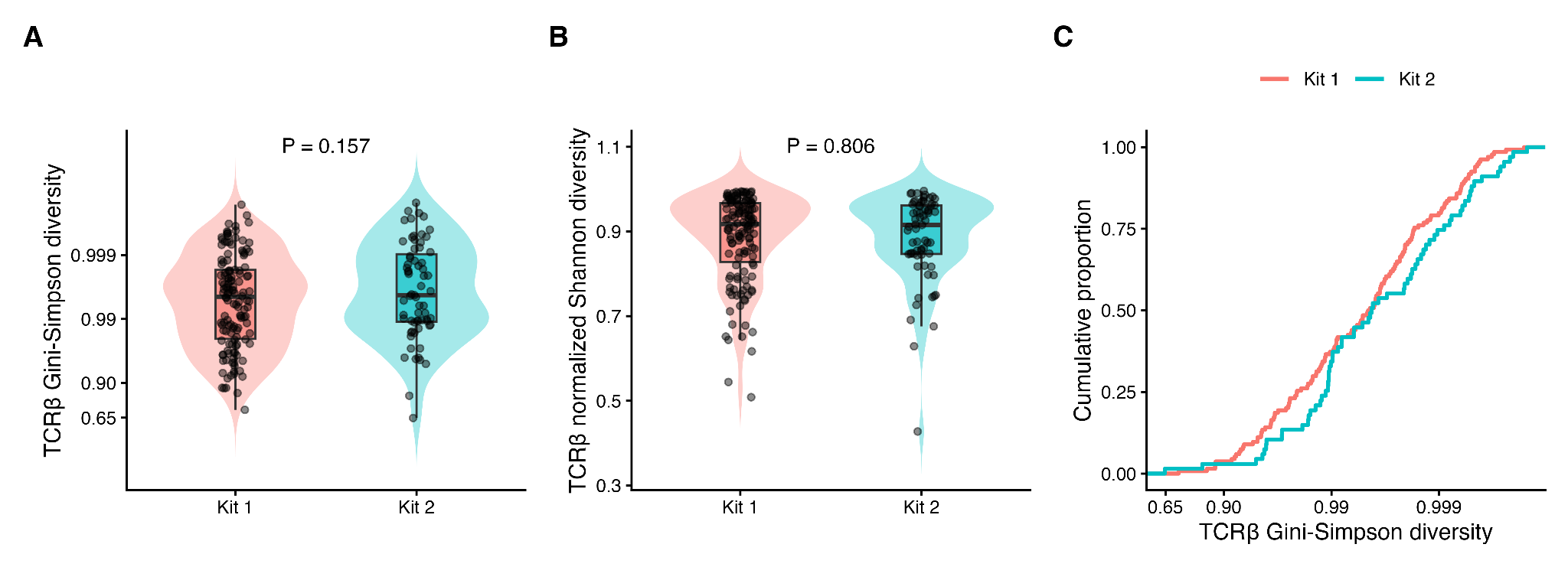

**Figure S9. Comparison of output from Cellecta DriverMap AIR TCR Profiling Kit versions 1 and 2.** (A) TCRβ Gini-Simpson diversity by kit. (B) TCRβ normalized Shannon diversity by kit. (C) Cumulative proportion of samples according to TCRβ Gini-Simpson diversity. Red = Version 1. Blue = Version 2. For visualization, TCRβ Gini-Simpson diversity was transformed as −log_10_(1 – Gini-Simpson) in panels A and C; axis labels show the original values. Differences between kit versions were assessed using the Wilcoxon rank-sum test on untransformed values. Abbreviations: TCRβ, T-cell receptor β.
